# Machine learning algorithm to characterize antimicrobial resistance associated with the International Space Station surface microbiome

**DOI:** 10.1101/2022.02.07.479455

**Authors:** Pedro Madrigal, Nitin K. Singh, Jason M. Wood, Elena Gaudioso, Félix Hernández-del-Olmo, Christopher E. Mason, Kasthuri Venkateswaran, Afshin Beheshti

## Abstract

**Background:** Antimicrobial Resistance (AMR) has a detrimental impact on human health on Earth and it is equally concerning in other environments such as space due to microgravity, radiation and confinement, especially for long-distance space travel. The International Space Station (ISS) is ideal for investigating microbial diversity and virulence. The shotgun metagenomics data of the ISS generated during the Microbial Tracking – 1 (MT-1) project and resulting metagenome-assembled genomes (MAGs) across three flights in eight different locations during 12 months were used in this study. The objective of this study was to identify the AMR genes associated with whole genomes of 227 cultivable strains, 21 shotgun metagenome sequences, and 24 MAGs retrieved from the ISS environmental samples that were treated with propidium monoazide (PMA; viable microbes).

**Results:** We have analyzed the data using a deep learning model, allowing us to go beyond traditional cut-offs based only on high DNA sequence similarity and extending the catalog of AMR genes. Our results in PMA treated samples revealed AMR dominance in the last flight for *Kalamiella piersonii*, a bacteria related to urinary tract infection in humans. The analysis of 227 pure strains isolated from the MT-1 project revealed hundreds of antibiotic resistance genes from many isolates, including two top-ranking species that corresponded to strains of *Enterobacter bugandensis* and *Bacillus cereus*. Computational predictions were experimentally validated by antibiotic resistance profiles in these two species, showing a high degree of concordance. Specifically, disc assay data confirmed the high resistance of these two pathogens to various beta-lactam antibiotics.

**Conclusion:** Overall, our computational predictions and validation analyses demonstrate the advantages of machine learning to uncover concealed AMR determinants in metagenomics datasets, expanding the understanding of the ISS environmental microbiomes and their pathogenic potential in humans.

## BACKGROUND

According to the World Health Organization, the widespread use of antibiotics worldwide and the slow discovery of major types on antibiotics in the last thirty years has made antibiotic resistance one of the biggest threats to human health, food security, and development (WHO, 2015). Accordingly, with NASA setting the course to return to the Moon with the Artemis mission and eventually venture out to Mars, maintaining the health of astronauts during long-term spaceflight is of paramount importance (Afshinnekoo et al., 2020). One area of particular concern is the reported increase in virulence and antibiotic resistance of microorganisms in space experiments (Juergensmeyer et al., 1999; Nickerson et al., 2004; Taylor, 2015; Zea et al., 2017; Wilson et al., 2017; Aunins et al., 2018; Urbaniak et al., 2018). Combined with a depressed or altered immune response in astronauts (Sonnenfeld and Shearer, 2002; Garrett-Bakelman et al., 2019), there is an increased risk of opportunistic microbial infection. Spaceflight promotes biofilm formation (Kim et al., 2013), and bacteria cultured from astronauts during flight were more resistant than isolates obtained from the same individual either pre- or post-flight (Tixador et al., 1985). Mutations also occurred more frequently in long-term spaceflights (Fukuda et al., 2000). An alternative non-mutually exclusive hypothesis to increased virulence or microbial resistance to antibiotics is that spaceflight conditions might alter the stability of pharmaceuticals (Du et al., 2011). In any case, bacterial infections might be more challenging to treat in space.

The International Space Station (ISS) is a closed-built environment with its own environmental microbiome shaped by microgravity, radiation, and limited human presence (Venkateswaran et al., 2014). We and others have shown that microbiomes are dynamic, diverse and sometimes intertwined at the ISS. Be NA et al. (2017) analyzed antibiotic resistance and virulence genes from dust and vacuum filter samples of ISS (treated with propidium monoazide, or PMA), demonstrating that human skin-associated microbes impact the ISS microbiome. Indeed, the skin and intestinal microbiomes of astronauts that spent 6 to 12 months in the ISS have been shown to be altered (Voorhies et al., 2019). In addition, the salivary microbiome of astronauts changed as a result of spaceflight, potentially activating microbes that promote viral replication (Urbaniak et al., 2020) and altering the abundance of some antimicrobial resistance (AMR) genes (Morrison et al., 2021). The ISS itself also presents specific core microbiome signatures on its surfaces that we characterized recently using shotgun metagenome and amplicon sequencing (Singh et al., 2018; Urbaniak et al., 2018; Checinska Sielaff et al., 2019), analogous to microbiome signatures found in specific geographies on Earth (Danko et al., 2021).

Further analyses across several missions have revealed that the microbiome of the crew’s skin resembled those of the surfaces inside the ISS collected by the crewmember on the same flight (Avila-Herrera et al., 2020). To better understand the composition of these bacterial populations we and others have characterized shotgun whole genome sequencing (WGS) of several ISS microorganisms (Singh et al., 2019; Bijlani et al., 2020; Bijlani et al., 2020b). Although most of them have been found to be non-pathogenic to humans, there are exceptions such as antibiotic-resistant *Enterobacter bugandensis* strains that could have an increased chance of pathogenicity (Singh et al., 2018b).

Computational analyses of microbiome data collected in Earth have shown that AMR can be predicted from genomic sequence of pure cultures alone (Hendriksen et al., 2019; Su et al., 2019), but a consensus approach on the best way to detect AMRs in metagenomic datasets has yet to be established (Ruppé et al., 2019). Generally, predictions are restricted to high identity (high sequence similarity to databases) cut-offs, requiring a ‘best-hit’ on an appropriate AMR database with a sequence identity greater than 80% by many programs such as ResFinder (Zankari et al., 2012). Although the ‘best-hit’ approach has a low false-positive rate, the false-negative rate can be very high, and a large number of actual Antibiotic Resistance Genes (ARGs) are predicted as non-ARGs, thus concealing the identification of potentially functional ARGs (Arango-Argoty et al., 2018). Another method of identification is to link the immune repertoire of the astronaut to the peptides of the microbes on the ISS, but this requires complex coordination with crew sampling and is rare (Danko et al., 2020). However, it has been shown recently that deep learning, a class of machine learning algorithms, can expand the catalog of AMR genes and increase the accuracy of the predictions based on metagenomic data (Arango-Argoty et al., 2018; Boolchandani et al., 2019; Hadjadj et al., 2019). We then hypothesized that the characterization of AMR from sequencing data at the ISS could be investigated from an artificial intelligence perspective using a robust deep learning framework. For that, we analyzed whole-genome sequences of 227 pure strains (cultivable microbes), metagenome sequences of 21 environmental samples, and 24 MAGs retrieved from PMA treated samples (**Fig. 1**).

**Figure 1.**
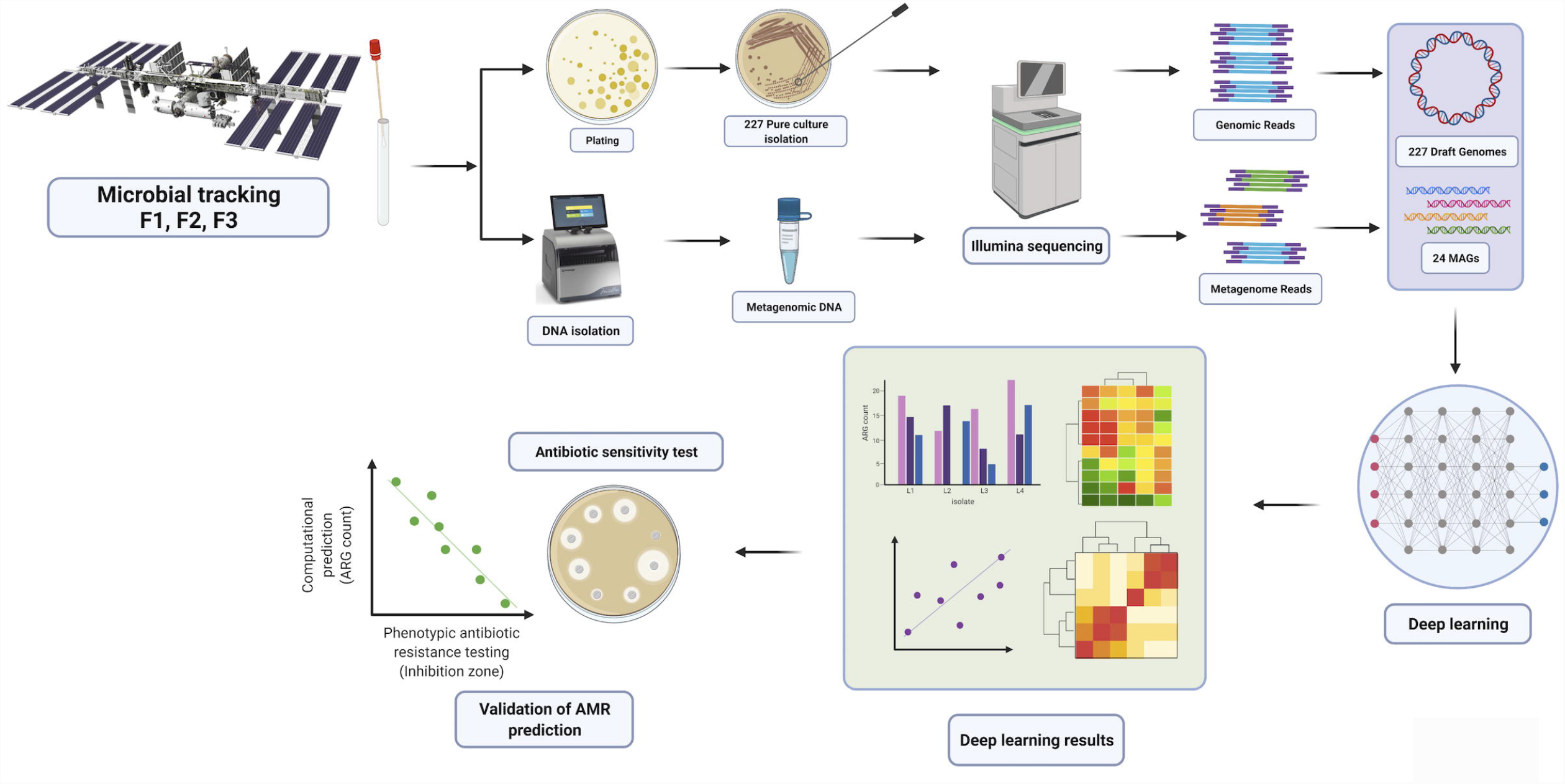
Overview of sample collection and data analysis for the characterization of antibiotic resistance at the ISS using deep learning. The data are processed in a step-wise fashion including data QC, mapping, quantification, and matching to time of collection and mission. The figure has been generated using BioRender (http://biorender.com).

## RESULTS

### Predictions based on short metagenomics sequences and ORFs partly overlap with previous analyses and reveal new AMR determinants at the ISS surface microbiome

The first shotgun metagenome sequences of intact microbial cells (Propidium monoazide-PMA treated) without whole genome amplification was performed by Singh et al. (2018). There, samples were taken in 8 locations across three flights (F1, F2, F3) during a period of 12 months. A detailed description of sampling procedures and locations can be found in Singh et al. (2018). To deploy a deep learning approach for predicting antibiotic resistance genes from metagenomic data, we used DeepARG, a computational resource proven to be more accurate than traditional approaches (Arango-Argoty et al., 2018). The model was trained using a merged database created after carefully curating three major databases (CARD, ARDB and UNIPROT). We first run DeepARG-SS (DeepARG for short reads) using the recommended prediction probability cut-off of 0.8 to obtain read counts of AMR genes (**Fig. 2a**). As in the seminal paper (Singh et al., 2018), quantification of antibiotics associated with AMR revealed ‘beta lactams’ ranking first and ‘peptide’ second, and generally more AMR reads counts observed in Flight 3 (F3) than in previous two flights (**Fig. 2a**). However, reads counts in certain antibiotics such as pleuromutilin, mupirocin and rifamycin were found largely in Flight 2 (**Fig. 2a**). Our read counts correlate (*r* = 0.86, *p* = 6.879e-7; Pearson’s product-moment correlation) with read counts obtained for antimicrobial resistance by Singh et al., (2018) (**Fig. 2b**). Taken together, these suggest a partial overlap with results obtained in Singh et al. (2018) analyzed using the traditional approach.

**Figure 2.**
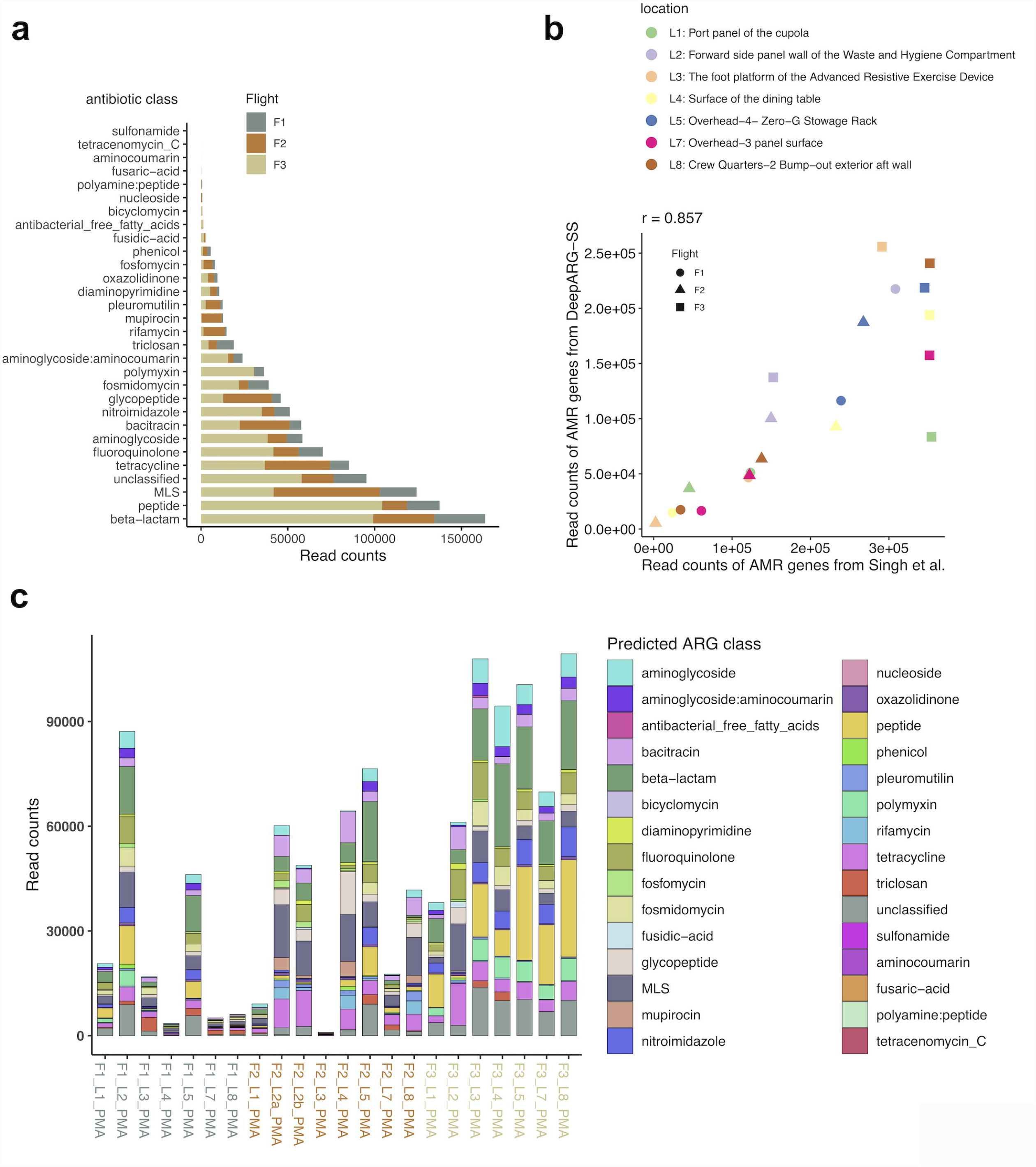
Prediction of ARGs using a pre-trained DeepARG-SS model. **(a)** Distribution of ARG read counts across antibiotic classes for the three flights (F1, F2, F3). **(b)** Correlation of read counts found by DeepARG-SS and those in Singh et al. (2018). Pearson’s product-moment correlation *r* = 0.86, (*p* = 6.879e-07) for the three flights and their locations. **(c)** Read counts of ARG class across flights for each location for PMA-treated samples in Singh et al. (2018). The antibiotic class (*multi-drug*) is not shown. Results are for ARGs with probability > 0.8.

While more AMR reads counts were found in Flight 3, we also observed variability between the different locations and flights, and an increasing number of read counts associated with time. For instance, location 4 (L4, surface of the dining table) increased the number of AMR reads counts with successive flights (**Fig. 2b-c**). While resistance to ‘beta lactams’ was evenly distributed across flights and locations, resistance to ‘polymyxin’ and specially ‘peptide’ represents a more significant proportion of AMR counts in locations of Flight 3 (**Fig. 2c**). In addition, we also observed the widespread presence of reads related to Macrolides, Lincosamides, Streptogamines (MLS), and tetracycline resistance.

To investigate the possible association between AMR patterns and specific microbes, we assembled the short reads into Metagenome-Assembled Genomes (MAGs; see Methods), identified their Open Reading Frames (ORFs), and repeated the prediction of ARGs using DeepARG-LS (Arango-Argoty et al., 2018). **Fig. 3a** shows the distribution of DeepARG classification probabilities and best-hit identity of ARGs in MAGs from the ISS. As we can retrieve highly probable ARGs (probability > 0.8) presenting low sequence identity (for many ARGs, identity is <40%), this method is likely more advantageous than using the ‘best-hit’ approach only. Compared to DeepARG-SS results obtained previously, the analysis of MAGs did not reveal significant differences in the number of ARGs predicted in the ORFs for the different flights (**Fig. 3b**). However, interestingly the results show a smaller number bacterial species having ARGs in Flight #1 (F1) when compared to Flights #2 and #3 (**Fig. 3b-c**) (data is shown for MAGs with at least 1 predicted ARG; the total number of MAGs analyzed is 24). Specifically, the number of locations is smaller in Flight 1 (3) than in F2 (n = 6) and F3 (n = 7) (**Fig. 3b**). Many ARGs were identified in *K. piersonii* MAGs in multiple locations during F3, showing AMR patterns related to (glyco)peptide, fluoroquinolone and MLS. (**Fig. 3c**). Of note, the *K. piersonii* strain closely related to one found at the ISS has been related to human urinary tract infection (Rheka et al., 2020). The potentially very pathogenic microbe *E. bugandensis* was found in location 2 (forward side panel wall of the Waste and Hygiene Compartment) in flight 1, presenting more than 40 ARGs. In addition, in the original study, *Pantoea* species were found to be the dominant genus in samples in 5 out of 7 locations sampled from Flight 3, especially at location 5 (surface rack). In our re-analysis, we observed *Pantoea brenneri* and *Pantoea dispersa* having ARGs related to beta-lactams and peptide (Singh et al., 2018), as well as to triclosan and polymyxin resistance.

**Figure 3.**
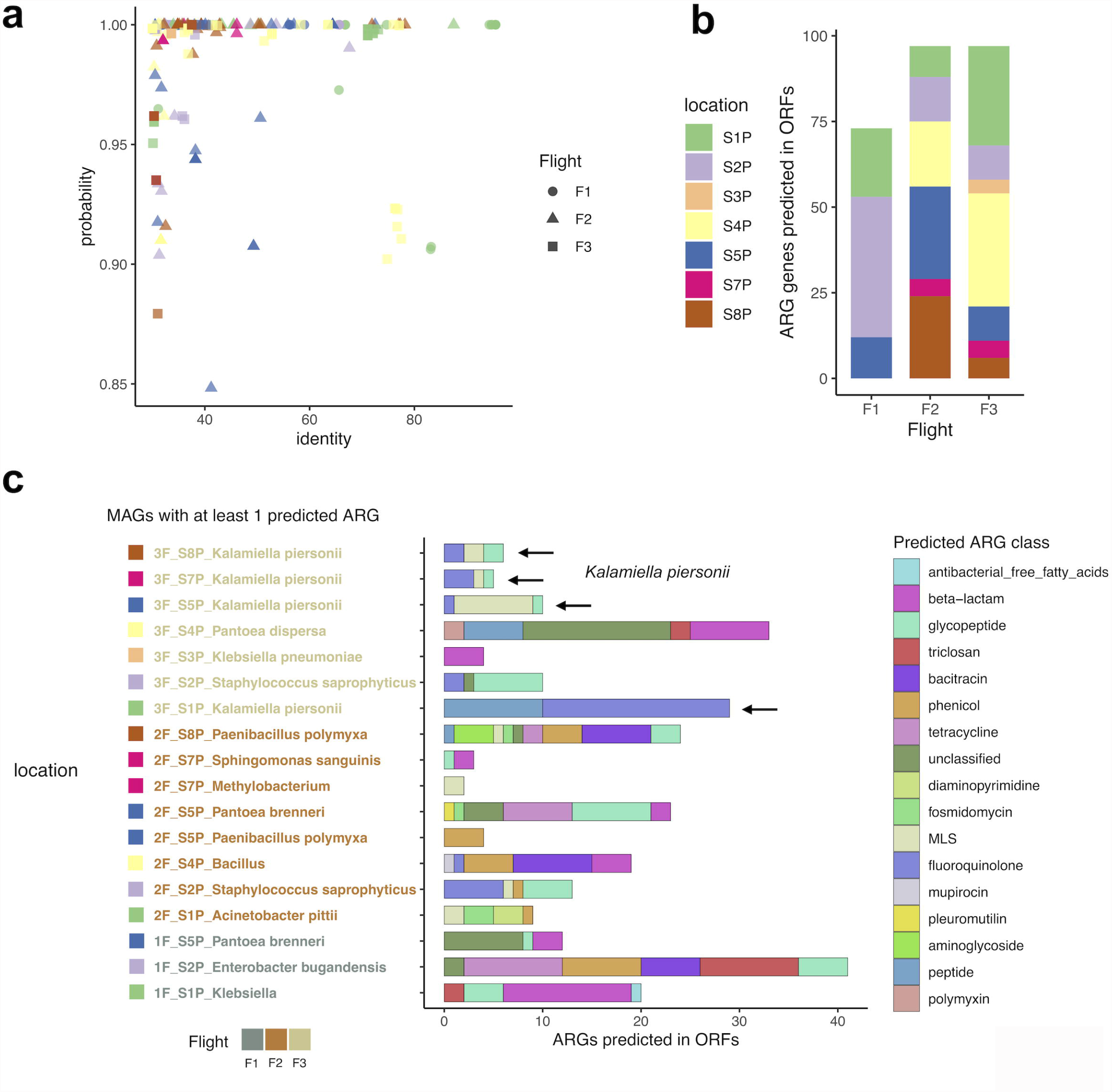
ARGs detected in ORFs in metagenome-assembled genomes (MAGs) from PMA-treated samples. **(a)** Distribution of DeepARG classification probability and best-hit identity in MAGs retrieved from the ISS. **(b)** Total number of ARGs predicted for each flight and location. **(c)** Number of ARGs precited for each MAG. Most common antibiotic class (*multi-drug*) not shown. The black arrows indicate *Kalamiella piersonii*.

Overall, by applying a deep learning approach, our results partially agree with earlier findings while providing new insights into previously unobserved antibiotic resistance classes (of the 30 antibiotic resistance categories included in the model). Specifically, the re-analysis of short sequences and MAGs from the ISS reveals dominance of *K. piersonii* antibiotic resistance in different locations of Flight 3 (**Fig. 3c**).

### Distribution of antibiotic resistance genes in scaffolds of Microbial Tracking-1 strains isolated from the ISS

We then applied DeepARG-LS to 227 Microbial Tracking-1 (MT-1) isolates (Mason and Venkateswaran labs, published and unpublished WGS of MT-1 pure strains isolated from ISS environment). We found a range of 2 to 92 ARGs in 184 out of 227 isolates (**Fig. 4a**; **Table 1**). Our machine learning approach allowed us to go beyond the traditional cut-off based only on high sequence DNA similarity (**Fig. S1**). These results suggest a widespread presence of potential ARGs in the isolates, with ‘multi-drug’ class being first, followed by glycopeptides, beta-lactams, bacitracin and tetracyclines. The ‘multi-drug’ antibiotic class was defined by aggregating several antibiotic names from the CARD and ARDB databases (efflux, multi-drug and na_antimicrobials). We then used BLAST to match isolates showing AMR sequences predicted by DeepARG to microbial species (**Fig. 4a**) and identified *Bacillus cereus* and *E. bugandensis*, which were previously profiled organisms on the ISS (Venkateswaran et al., 2017; Singh et al., 2018b) as the top 2 ranking species with a high number of ARGs. We have previously shown that five *E. bugandensis* isolates were almost equivalent to nosocomial earth isolates showing resistance to multi-drug antibiotic compounds, fluoroquinolones, and fosfomycin (Singh et al., 2018b). In addition, *E. bugandensis* strains were shown to be resistant to 9 antibiotics (Urbaniak et al., 2018). Our results reinforce the potential pathogenicity of this microbe. Nonetheless, antimicrobial resistance was not examined for *B. cereus* strains in Venkateswaran et al. (2017). *B. cereus* is a food poisoning microorganism that might be a concern for crew members’ health. In addition, we found novel ARGs associated with other species such as *K. pneumoniae, Pantoea, Paenibacillus polymyxa, B. velezensis, E. faecalis, Sphingomonas*, and, with a lower number of ARGs, several species of *Staphylococcus. E. faecalis* virulence was previously shown to be affected by microgravity (Hammond et al., 2013).

**Figure 4.**
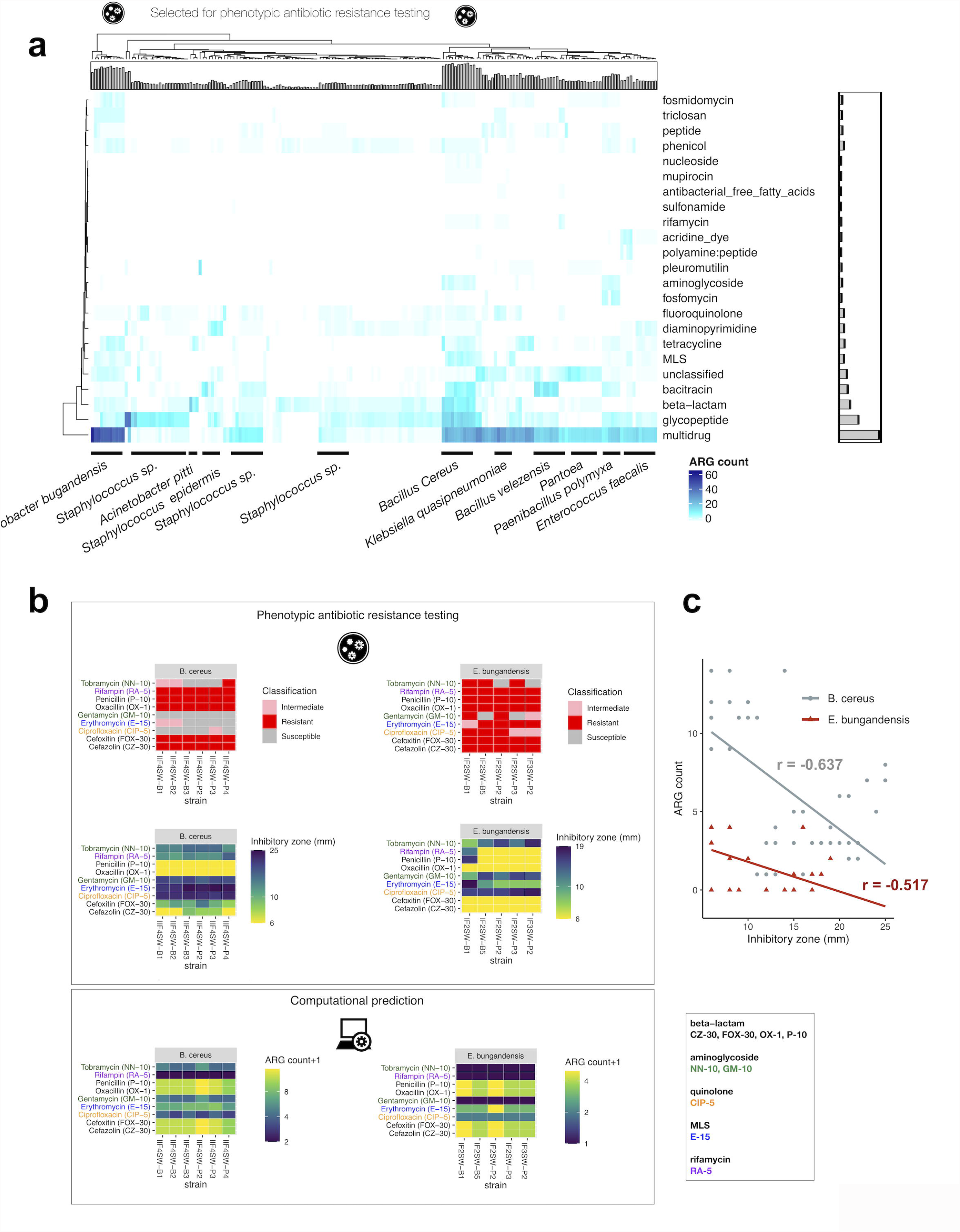
Heatmap and clustering of ARG counts detected in MT-1 pure strains isolated from the ISS and AST validations. **(a)** Heatmap with ARG count. The barplots illustrate the number of ARGs across rows and across columns. Species were identified using BLAST. Only ARGs with probability > 0.8 were considered, as recommended. **(b)** Antibacterial Susceptibility Tests (AST) on *E. bugandensis* and *B cereus* strains for several antibiotics (top), and comparison with machine learning predictions shown in Fig. 4a (bottom). **(c)** Scatterplot of zone of inhibition value (in mm.) and ARG count shown in (b), together with a linear model fit. Pearson’s product-moment correlation values are indicated.

To experimentally validate machine learning predictions on previously unobserved AMR patterns above, we performed Antibacterial Susceptibility Tests (AST) for the species found to be potentially most pathogenic, in our case *E. bugandensis* and *B cereus* as they have a higher number of ARGs (**Table S1; Fig. 4a**). For that, we use disc diffusion on strains isolated at the ISS for the following antimicrobials: Cefazolin (beta− lactam), Cefoxitin (beta− lactam), Ciprofloxacin (quinolone), Erythromycin (MLS), Gentamycin (aminoglycoside), Oxacillin (beta− lactam), Penicillin (beta− lactam), Rifampin (rifamycin) and Tobramycin (aminoglycoside) (**Fig. 4b**). The prediction patterns closely matched the AST results (**Fig. 4b**), although DeepARG failed to detect Rifampin resistance, especially for *E. bugandensis*.

Although different antibiotics have different inhibitory zone cut-offs for a strain to be considered as resistant (**Table S2**), remarkably we found an inverse correlation between the zone of inhibition and ARG count for *B. cereus* (*r*= -0.637, Pearson’s product-moment correlation, *p*= 2.2e-7) and *E. bugandensis* (*r*= -0.517; *p*= 0.0002765) (**Fig. 4c**), demonstrating the applicability and high accuracy of computational prediction of AMR for microbiome data obtained in space.

## DISCUSSION

Many ARGs that present high probability but low sequence identity to known sequences will be missed using traditional ‘best-hit’ approaches that require a high degree of sequence identity. To solve this, computational methods have been developed to identify AMR in genomes and metagenomes (Arango-Argoty et al., 2018; Berglund et al., 2019; Chowdhury et al., 2019; Lakin et al., 2019; Ruppé et al., 2019). Despite these developments, a consensus approach to detect AMR in metagenomics datasets is yet to be defined (Ruppé et al., 2019). The objective of this study was to identify the AMR genes associated with cultivated strains and metagenomes generated from the ISS environmental surfaces using an accurate deep learning approach (**Fig. 1**).

Firstly, we re-analyzed shotgun metagenome sequences of 21 environmental samples that were treated with PMA (viable microbes), and their associated 24 metagenome-assembled genomes (MAG) retrieved from the PMA-treated samples. The re-analysis showed increased read counts associated with AMR and in more locations when considering MAGs, in flight 3 (**Fig. 2**). This could be explained due to the ISS crew being replaced during Flight 3. The abundance of *Enterobacteriaceae* in Flight 3 was discussed in Singh et al., (2018b). In addition, *K. piersonii* spread across four different locations (L1, L5, L7, L8) at Flight 3, presenting resistance to specific antibiotics (glyco/peptide, fluoroquinolone and MLS) (**Fig. 3c**). We have previously isolated strains from Locations 1, 2, 5, 6, and 7, defining a novel bacterial genus from the ISS samples (Singh et al., 2019). While *K. piersonii* do have virulence genes in the genome, a dichotomy was found as disc diffusion tests revealed multi-drug resistance, while the PathogenFinder algorithm predicted

*K. piersonii* strains as non-human pathogens. All seven *K. piersonii* isolates were resistant to cefoxitin (beta_lactam class in DeepARG), erythromycin (MLS), oxacillin (beta_lactam), penicillin (beta_lactam), and rifampin. At the same time, all strains were susceptible to cefazolin, ciprofloxacin (quinolone), and tobramycin (aminoglycoside) (Singh et al., 2019). The DeepARG database does not include some of these antibiotics, but we found AMR sequences related to resistance to (glyco)peptide, fluoroquinolone, and MLS, validating some previous results. Therefore, PathogenFinder (Cosentino et al., 2013) results in Singh et al. (2019) suggesting *K. piersonii* as a non-human pathogen should be treated with caution. Furthermore, the strain YU22 (closest match is IIIF1SW-P2^T^ detected as ISS) isolated in urine microbiome of a kidney stone patient has shown to be an uropathogenic bacteria, showing many virulence factors that are needed for host cell invasion and colonization (Rheka et al., 2020).

Secondly, the whole genome sequences (WGS) of 227 pure strains (cultivable microbes) were analyzed to identify AMR genes (**Fig. 4a**). We found the human pathogens *E. bugandensis* and *B. cereus* presenting many potential ARGs in the MT-1 scaffolds. Up to five strains isolated from the ISS have been closely related to the type strain EB-247^T^ and two clinical isolates (153_ECLO and MBRL 1077) and share similar AMR patterns (Singh et al., 2018b). 112 genes were found to be involved in virulence, disease, and defence in the ISS strains (Singh et al., 2018b). Our re-analysis confirms the *multi-drug* resistance to antibiotics for the ISS isolates, which is the highest among all the isolates. Unlike in Singh et al. (2018b), we found fluoroquinolone resistance low, and null for fosfomycin. Conversely, *B. cereus* is a gram-positive bacterium commonly found in food. After infection, most emetic patients recover within 24 hours, but in some cases, the toxin can be fatal via a fulminant hepatic failure (Mahler et al., 1997; Dierick et al., 2005). Overall, *multi-drug* resistance was found widespread in many microbes. Third, phenotypic antibiotic resistance testing data obtained from traditional antibiotic tests generated for biosafety level 2 (BSL-2) strains were compared with the computational approaches that predicted the presence of the AMR genes, showing an excellent agreement for the antibiotics tested (**Fig. 4b-c**). A disadvantage of the deep learning model used is that the prediction can disentangle the family of antibiotics but not specific compounds.

Many studies have shown the association between several microorganisms (bacterial, as well as phage and non-phase viral sequences) and several cancer features. Although it is unclear whether this corresponds to correlation or causation, the microbiome can undoubtedly be used as a cancer biomarker. For instance, certain strains of *Fusobacterium spp*. can be utilized as an independent diagnostic assay for colon cancer (Zhang et al., 2019). Therefore, a better understanding of the microbial communities and their degree of pathogenicity in surface-human microbiomes in space could also be useful for human health monitorization with detection and prognostic values in long term space travel. We are currently collecting more data for Microbial Tracking-2 (MT-2) and MT-3 missions. We plan to extend the AMR catalog, characterize microbe diversity, and monitor the evolution of AMR in longer time periods to discover new factors involved in pathogenicity of microorganisms exposed to space conditions.

## METHODS

### Metagenome-Assembled Genomes (MAGs) methodology

The paired-end 100-bp metagenomic reads from NCBI Short Read Archive (SRA) under the bio-project number PRJNA438545 were processed with Trimmomatic (Bolger et al., 2014) to trim adapter sequences and low-quality ends, with a minimum Phred score of 20 across the entire length of the read used as a quality cut-off. Reads shorter than 80-bp were removed after trimming. Remaining high-quality reads were subsequently assembled using metaSPAdes (Nurk et al., 2017). Contigs were binned using Metabat2 v2.11.3 (Kang et al., 2015). Recovered genomes were evaluated with CheckM (Parks et al., 2015), and a recovered genome was considered good with at least 90% completeness and at most 10% contamination. Each genome was subsequently annotated with the help of Rapid Annotations using Subsystems Technology (RAST), and near identifications were predicted (Aziz et al., 2008).

### Isolates from Microbial Tracking 1 mission

To create the whole-genome sequences (WGS) of these strains, shotgun libraries were prepared using the Illumina Nextera Flex protocol (Singh et al., 2018b), using NovaSeq 6000 S4 flow cell 2150 paired-end (PE) sequencing. Verification of the quality of the raw sequencing data was carried out using FastQC v0.11.7 (Andrews, 2015). Quality control for adapter trimming and quality filtering were performed using fastp v0.20.0 (Chen et al., 2018), and then SPAdes v3.11.1 (Bankevich et al., 2012) was used to assemble all the cleaned sequences. Fastp quality control was based on the following three parameters: (i) correction of mismatches in overlapped regions of paired-end reads, (ii) trimming of autodetected adapter sequences, and (iii) quality trimming at the 59 and 39 ends. To determine the quality of the assembled sequences, the number of contigs, the N50 value, and the total length were calculated using QUAST v5.0.2 (Gurevich et al., 2013). Default parameters were used for all software. The average nucleotide identity (ANI) (Yoon et al., 2017) was calculated using OrthoANIu by comparing each of the scaffolds to the WGS of the respective type strains.

### Identification of ORFs in microbial DNA sequences

Glimmer (Gene Locator and Interpolated Markov ModelER) v3.02 was used with default parameters to identify the coding regions and distinguish them from non-coding DNA in MAGs and MT-1 scaffolds that could be used as an input in DeepARG-LS. Minimum gene length was indicated as 50 bp (‘glimmer3 -g50’). Glimmer reads DNA sequences in a FASTA file format and predicts genes in them using an Interpolated Context Model (Delcher et al., 2007).

### Prediction of antibiotic resistance genes in short reads and full-gene length sequences

DeepARG version 2 (Arango-Argoty et al., 2018), a deep learning-based approach for predicting Antibiotic Resistance Genes (ARGs) and annotation, was run with the ‘--reads’ option (DeepARG-SS) for NGS reads and the ‘--genes’ option (DeepRG-LS) for longer gene-like sequences obtained with Glimmer. The DeepARG model consists of four dense hidden layers of 2000, 1000, 500, and 100 units that propagate a bit score distribution. The output layer of the deep neural network consists of 30 units that correspond to the antibiotic resistance categories (102 antibiotics consolidated into 30 antibiotic categories). The model was trained with a curated database of 14,933 genes from three databases (CARD, ARDB, and UNIPROT) (Arango-Argoty et al., 2018). Default options were used: 50% minimum percentage of identity to consider, significance of the prediction probability cut-off of 0.8 as recommended (Arango-Argoty et al., 2018), and E-value of alignments (default 1e-10). The software was downloaded from https://bitbucket.org/gusphdproj/deeparg-ss.

### Microbial Nucleotide BLAST

Nucleotide-Nucleotide BLAST 2.10.1+ (https://blast.ncbi.nlm.nih.gov/Blast.cgi) was used to identify microbial species associated to MT-1 scaffolds. Sequences producing significant alignments were ranked and the species associated to maximum Score (bits) and minimum E-value was deemed as the closest match.

### Phenotypic antibiotic resistance testing

Disc assays experiments were performed as in Urbaniak et al., (2018). The isolates were streaked from glycerol stocks onto R2A plates. A single colony was inoculated into 5 mL Tryptic Soy Broth (TSB) and grown overnight at 30°C. Aliquots of 100 µL were plated on TSA. Agar diffusion discs (BD BBLTM Sensi-DiscTM, Franklin Lakes, NJ) were placed aseptically on a plate and the strains were incubated at 37°C for 24 h. The tested antibiotics included: 30-µg cefazolin (CZ-30); 30-µg cefoxitin (FOX-30), 5-µg ciprofloxacin (CIP-5), 15 µg erythromycin (E-15), 10-µg gentamicin (GM-10), 1-µg oxacillin (OX), 10-µg penicillin (P-10), 5-µg rifampin (RA-5), and 10-µg tobramycin (NN-10). The diameter of inhibition zones was measured for each antibiotic disk and recorded in millimeters. The resistance results were compared with the zone diameter interpretive charts provided by the manufacturer. When the spontaneous mutants were present in response to some antibiotics, they were isolated, subcultured and tested for the specific antibiotic resistance.

## Data availability

Raw metagenomics reads from three flights on multiple locations were downloaded from NASA GeneLab repository https://genelab.nasa.gov (GLDS-69). Microbial tracking-1 (MT-1) datasets were obtained from GLDS-67, GLDS-302, GLDS-303, GLDS-309, GLDS-311 and GLDS-350. The rest of the samples are deposited at DDBJ/ ENA/GenBank or are unpublished.

## SUPPLEMENTARY INFORMATION

**Additional file 1: Figure S1**. Distribution of DeepARG classification probability and best-hit identity in MT-1 pure strains isolated from the ISS. The blue dashed line indicates 50% sequence identity.

**Additional file 2: Table S1**. Rank of MT-1 isolates, ordered by number of ARGs predicted, shown in Figure 4a. Species information obtained from Microbial Nucleotide BLAST.

**Additional file 3: Table S2**. Phenotypic antibiotic resistance testing results for *E. bugandensis* and *B cereus*.

**Additional file 4: File S1**. Raw results of the DeepARG analyses (zip compressed).

## Acknowledgments

We thank Dr Sylvain Costes, Dr Jonathan Galazka and Dr Daniel C. Berrios for the initial conversations that inspired this project. The authors would like to acknowledge the members of the Microbiome and Multi-Omics/Systems Biology Analysis Working Groups of NASA GeneLab. Part of the research described in this manuscript was performed at the Jet Propulsion Laboratory, California Institute of Technology under a contract with NASA. We would like to thank Microbial Tracking -1 and 2 members for isolating the strain and generating draft assembly of genomes. We thank Biotechnology and Planetary Protection Group members for supporting sample analyses. We also acknowledge the Jet Propulsion Laboratory supercomputing facility staff, notably Narendra J. Patel (Jimmy) and Edward Villanueva, for their continuous support in providing the best possible infrastructure for BIG-DATA analysis.

## Authors’ contributions

PM performed the computational analyses and led the writing. NS, KV, JW, and CEM provided data, exchange of ideas, and edits to the manuscript. EG and FHdO contributed to machine learning aspects. AB supervised and supported the study, wrote and provided final approval of the manuscript. All authors provided feedback and contributed to the research and final manuscript.

## Availability of data and materials

The code use for the analysis is available at https://github.com/pmb59/AMRISS.

## DECLARATIONS

### Ethics approval and consent to participate

Not applicable.

### Consent for publication

All authors have read and agreed with the manuscript.

## REFERENCES

Afshinnekoo E, Scott RT, MacKay MJ, Pariset E, Cekanaviciute E, Barker R, Gilroy S, Hassane D, Smith SM, Zwart SR, Nelman-Gonzalez M, Crucian BE, Ponomarev SA, Orlov OI, Shiba D, Muratani M, Yamamoto M, Richards SE, Vaishampayan PA, Meydan C, Foox J, Myrrhe J, Istasse E, Singh N, Venkateswaran K, Keune JA, Ray HE, Basner M, Miller J, Vitaterna MH, Taylor DM, Wallace D, Rubins K, Bailey SM, Grabham P, Costes SV, Mason CE, Beheshti A. Fundamental Biological Features of Spaceflight: Advancing the Field to Enable Deep-Space Exploration. Cell. 2020 Nov 25;183(5):1162–1184. doi: 10.1016/j.cell.2020.10.050

Andrews S. 2015. FastQC: a quality tool for high throughput sequence data. http://www.bioinformatics.babraham.ac.uk/projects/fastqc/.

Arango-Argoty G, Garner E, Pruden A, Heath LS, Vikesland P, Zhang L. DeepARG: a deep learning approach for predicting antibiotic resistance genes from metagenomic data. Microbiome. 2018 Feb 1;6(1):23. doi: 10.1186/s40168-018-0401-z.

Aunins TR, Erickson KE, Prasad N, et al. Spaceflight Modifies Escherichia coli Gene Expression in Response to Antibiotic Exposure and Reveals Role of Oxidative Stress Response. Front Microbiol. 2018;9:310. Published 2018 Mar 16. doi:10.3389/fmicb.2018.00310

Avila-Herrera A, Thissen J, Urbaniak C, et al. Crewmember microbiome may influence microbial composition of ISS habitable surfaces. PLoS One. 2020;15(4):e0231838

Aziz, R.K., Bartels, D., Best, A.A., Dejongh, M., Disz, T., Edwards, R.A., Formsma, K., Gerdes, S., Glass, E.M., Kubal, M., Meyer, F., Olsen, G.J., Olson, R., Osterman, A.L., Overbeek, R.A., Mcneil, L.K., Paarmann, D., Paczian, T., Parrello, B., Pusch, G.D., Reich, C., Stevens, R., Vassieva, O., Vonstein, V., Wilke, A., and Zagnitko, O. (2008). The RAST Server: rapid annotations using subsystems technology. BMC Genomics 9, 75.

Bankevich A, Nurk S, Antipov D, Gurevich AA, Dvorkin M, Kulikov AS, Lesin VM, Nikolenko SI, Pham S, Prjibelski AD, Pyshkin AV, Sirotkin AV, Vyahhi N, Tesler G, Alekseyev MA, Pevzner PA. 2012. SPAdes: a new genome assembly algorithm and its applications to single-cell sequencing. J Comput Biol 19:455–477. https://doi.org/10.1089/cmb.2012.0021.

Be NA, Avila-Herrera A, Allen JE, et al. Whole metagenome profiles of particulates collected from the International Space Station. Microbiome. 2017;5(1):81. Published 2017 Jul 17. doi:10.1186/s40168-017-0292-4

Berglund F, Österlund T, Boulund F, Marathe NP, Larsson DGJ, Kristiansson E. Identification and reconstruction of novel antibiotic resistance genes from metagenomes. Microbiome. 2019;7(1):52. Published 2019 Apr 1. doi:10.1186/s40168-019-0670-1

Bijlani S, Singh NK, Mason CE, Wang CCC, Venkateswaran K. Draft Genome Sequences of Sphingomonas Species Associated with the International Space Station. Microbiol Resour Announc. 2020;9(25):e00578–20. Published 2020 Jun 18. doi:10.1128/MRA.00578-20

Bijlani S, Singh NK, Mason CE, Wang CCC, Venkateswaran K. Draft Genome Sequences of Tremellomycetes Strains Isolated from the International Space Station. Microbiol Resour Announc. 2020b;9(26):e00504–20. Published 2020 Jun 25. doi:10.1128/MRA.00504-20

Bolger, A.M., Lohse, M., and Usadel, B. (2014). Trimmomatic: a flexible trimmer for Illumina sequence data. Bioinformatics 30, 2114–2120.

Boolchandani M, D’Souza AW, Dantas G. Sequencing-based methods and resources to study antimicrobial resistance. Nat Rev Genet. 2019;20(6):356–370. doi:10.1038/s41576-019-0108-4

Checinska Sielaff A, Urbaniak C, Mohan GBM, Stepanov VG, Tran Q, Wood JM, Minich J, McDonald D, Mayer T, Knight R, Karouia F, Fox GE, Venkateswaran K. Characterization of the total and viable bacterial and fungal communities associated with the International Space Station surfaces. Microbiome. 2019 Apr 8;7(1):50. doi: 10.1186/s40168-019-0666-x

Chen S, Zhou Y, Chen Y, Gu J. 2018. fastp: an ultra-fast all-in-one FASTQ preprocessor. Bioinformatics 34:i884–i890. https://doi.org/10.1093/bioinformatics/bty560.

Chowdhury AS, Call DR, Broschat SL. Antimicrobial Resistance Prediction for Gram-Negative Bacteria via Game Theory-Based Feature Evaluation [published correction appears in Sci Rep. 2020 Jan 30;10(1):1846]. Sci Rep. 2019;9(1):14487. Published 2019 Oct 9. doi:10.1038/s41598-019-50686-z

Cosentino S, Voldby Larsen M, Møller Aarestrup F, Lund O. PathogenFinder--distinguishing friend from foe using bacterial whole genome sequence data [published correction appears in PLoS One. 2013;8(12). doi:10.1371/annotation/b84e1af7-c127-45c3-be22-76abd977600f]. PLoS One. 2013;8(10):e77302. Published 2013 Oct 28. doi:10.1371/journal.pone.0077302

Danko D, Bezdan D, Afshin EE, Ahsanuddin S, Bhattacharya C, Butler DJ, Chng KR, Donnellan D, Hecht J, Jackson K, Kuchin K, Karasikov M, Lyons A, Mak L, Meleshko D, Mustafa H, Mutai B, Neches RY, Ng A, Nikolayeva O, Nikolayeva T, Png E, Ryon KA, Sanchez JL, Shaaban H, Sierra MA, Thomas D, Young B, Abudayyeh OO, Alicea J, Bhattacharyya M, Blekhman R, Castro-Nallar E, Cañas AM, Chatziefthimiou AD, Crawford RW, De Filippis F, Deng Y, Desnues C, Dias-Neto E, Dybwad M, Elhaik E, Ercolini D, Frolova A, Gankin D, Gootenberg JS, Graf AB, Green DC, Hajirasouliha I, Hastings JJA, Hernandez M, Iraola G, Jang S, Kahles A, Kelly FJ, Knights K, Kyrpides NC, Łabaj PP, Lee PKH, Leung MHY, Ljungdahl PO, Mason-Buck G, McGrath K, Meydan C, Mongodin EF, Moraes MO, Nagarajan N, Nieto-Caballero M, Noushmehr H, Oliveira M, Ossowski S, Osuolale OO, Özcan O, Paez-Espino D, Rascovan N, Richard H, Rätsch G, Schriml LM, Semmler T, Sezerman OU, Shi L, Shi T, Siam R, Song LH, Suzuki H, Court DS, Tighe SW, Tong X, Udekwu KI, Ugalde JA, Valentine B, Vassilev DI, Vayndorf EM, Velavan TP, Wu J, Zambrano MM, Zhu J, Zhu S, Mason CE; International MetaSUB Consortium. A global metagenomic map of urban microbiomes and antimicrobial resistance. Cell. 2021 184(13):3376–3393

Danko DC, Singh N, Butler DJ, Mozsary C, Jiang P, Keshavarzian A, Maienschein-Cline M, Chlipala G, Afshinnekoo E, Bezdan D, Garrett-Bakelman F, Green SJ, Turek FW, Vitaterna MH, Venkateswaran K, Mason CE. Genetic and Immunological Evidence for Microbial Transfer Between the International Space Station and an Astronaut. bioRxiv 2020.11.10.376954; doi: https://doi.org/10.1101/2020.11.10.376954

Delcher AL, Bratke KA, Powers EC, Salzberg SL. Identifying bacterial genes and endosymbiont DNA with Glimmer. Bioinformatics. 2007;23(6):673–679. doi:10.1093/bioinformatics/btm009

Dierick, Katelijne; Van Coillie, Els; Swiecicka, Izabela; Meyfroidt, Geert; et al. (August 2005). “Fatal family outbreak of Bacillus cereus-associated food poisoning”. Journal of Clinical Microbiology. 43 (8): 4277–4279.

Du B, Daniels VR, Vaksman Z, Boyd JL, Crady C, Putcha L. Evaluation of physical and chemical changes in pharmaceuticals flown on space missions. AAPS J. 2011;13(2):299–308. doi:10.1208/s12248-011-9270-0

Fukuda T, Fukuda K, Takahashi A, et al. Analysis of deletion mutations of the rpsL gene in the yeast Saccharomyces cerevisiae detected after long-term flight on the Russian space station Mir. Mutat Res. 2000;470(2):125–132. doi:10.1016/s1383-5742(00)00054-5

Garrett-Bakelman FE, Darshi M, Green SJ, Gur RC, Lin L, Macias BR, McKenna MJ, Meydan C, Mishra T, Nasrini J, Piening BD, Rizzardi LF, Sharma K, Siamwala JH, Taylor L, Vitaterna MH, Afkarian M, Afshinnekoo E, Ahadi S, Ambati A, Arya M, Bezdan D, Callahan CM, Chen S, Choi AMK, Chlipala GE, Contrepois K, Covington M, Crucian BE, De Vivo I, Dinges DF, Ebert DJ, Feinberg JI, Gandara JA, George KA, Goutsias J, Grills GS, Hargens AR, Heer M, Hillary RP, Hoofnagle AN, Hook VYH, Jenkinson G, Jiang P, Keshavarzian A, Laurie SS, Lee-McMullen B, Lumpkins SB, MacKay M, Maienschein-Cline MG, Melnick AM, Moore TM, Nakahira K, Patel HH, Pietrzyk R, Rao V, Saito R, Salins DN, Schilling JM, Sears DD, Sheridan CK, Stenger MB, Tryggvadottir R, Urban AE, Vaisar T, Van Espen B, Zhang J, Ziegler MG, Zwart SR, Charles JB, Kundrot CE, Scott GBI, Bailey SM, Basner M, Feinberg AP, Lee SMC, Mason CE, Mignot E, Rana BK, Smith SM, Snyder MP, Turek FW. The NASA Twins Study: A multidimensional analysis of a year-long human spaceflight. Science. 2019 Apr 12;364(6436):eaau8650. doi: 10.1126/science.aau8650

Gurevich A, Saveliev V, Vyahhi N, Tesler G. 2013. QUAST: quality assessment tool for genome assemblies. Bioinformatics 29:1072–1075. https://doi.org/10.1093/bioinformatics/btt086.

Hadjadj L, Baron SA, Diene SM, Rolain JM. How to discover new antibiotic resistance genes?. Expert Rev Mol Diagn. 2019;19(4):349–362. doi:10.1080/14737159.2019.1592678

Hammond TG, Stodieck L, Birdsall HH, et al. Effects of microgravity on the virulence of Listeria monocytogenes, Enterococcus faecalis, Candida albicans, and methicillin-resistant Staphylococcus aureus. Astrobiology. 2013;13(11):1081–1090. doi:10.1089/ast.2013.0986

Hendriksen RS, Bortolaia V, Tate H, Tyson GH, Aarestrup FM, McDermott PF. Using Genomics to Track Global Antimicrobial Resistance. Front Public Health. 2019;7:242. Published 2019 Sep 4. doi:10.3389/fpubh.2019.00242

Juergensmeyer MA, Juergensmeyer EA, Guikema JA. Long-term exposure to spaceflight conditions affects bacterial response to antibiotics. Microgravity Sci Technol. 1999;12(1):41

Kang, D.D., Froula, J., Egan, R., and Wang, Z. (2015). MetaBAT, an efficient tool for accurately reconstructing single genomes from complex microbial communities. PeerJ 3, e1165.

Kim W, Tengra FK, Young Z, et al. Spaceflight promotes biofilm formation by Pseudomonas aeruginosa. PLoS One. 2013;8(4):e62437. Published 2013 Apr 29.

Lakin SM, Kuhnle A, Alipanahi B, et al. Hierarchical Hidden Markov models enable accurate and diverse detection of antimicrobial resistance sequences. Commun Biol. 2019;2:294. Published 2019 Aug 6. doi:10.1038/s42003-019-0545-9

Mahler, Hellmut; Pasi, Aurelio; Kramer, John M.; Schulte, Petra; et al. (17 April 1997). “Fulminant liver failure in association with the emetic toxin of Bacillus cereus”. The New England Journal of Medicine. 336 (16): 1142–1148.

Morrison MD, Thissen JB, Karouia F, Mehta S, Urbaniak C, Venkateswaran K, Smith DJ and Jaing C (2021) Investigation of Spaceflight Induced Changes to Astronaut Microbiomes. Front. Microbiol. 12:659179. doi: 10.3389/fmicb.2021.659179

Nickerson CA, Ott CM, Wilson JW, Ramamurthy R, Pierson DL. Microbial responses to microgravity and other low-shear environments. Microbiol Mol Biol Rev. 2004;68(2):345–361. doi:10.1128/MMBR.68.2.345-361.2004

Nurk, S., Meleshko, D., Korobeynikov, A., and Pevzner, P.A. (2017). metaSPAdes: a new versatile metagenomic assembler. Genome Res 27, 824–834.

Parks, D.H., Imelfort, M., Skennerton, C.T., Hugenholtz, P., and Tyson, G.W. (2015). CheckM: assessing the quality of microbial genomes recovered from isolates, single cells, and metagenomes. Genome Res 25, 1043–1055.

Rekha PD, Hameed A, Manzoor MAP, Suryavanshi MV, Ghate SD, Arun AB, Rao SS, Athmika, Bajire SK, Mujeeburahiman M, Young CC. First Report of Pathogenic Bacterium Kalamiella piersonii Isolated from Urine of a Kidney Stone Patient: Draft Genome and Evidence for Role in Struvite Crystallization. Pathogens. 2020 Aug 29;9(9):711. doi: 10.3390/pathogens9090711. PMID: 32872396; PMCID: PMC7558591.

Ruppé E, Ghozlane A, Tap J, et al. Prediction of the intestinal resistome by a three-dimensional structure-based method. Nat Microbiol. 2019;4(1):112–123. doi:10.1038/s41564-018-0292-6

Singh NK, Wood JM, Karouia F, Venkateswaran K. Succession and persistence of microbial communities and antimicrobial resistance genes associated with International Space Station environmental surfaces. Microbiome. 2018 Nov 13;6(1):204. doi: 10.1186/s40168-018-0585-2.

Singh NK, Bezdan D, Checinska Sielaff A, Wheeler K, Mason CE, Venkateswaran K. Multi-drug resistant Enterobacter bugandensis species isolated from the International Space Station and comparative genomic analyses with human pathogenic strains. BMC Microbiol 18:175 2018b

Singh NK, Wood JM, Mhatre SS, Venkateswaran K. Metagenome to phenome approach enables isolation and genomics characterization of Kalamiella piersonii gen. nov., sp. nov. from the International Space Station. Appl Microbiol Biotechnol. 2019;103(11):4483–4497. doi:10.1007/s00253-019-09813-z

Sonnenfeld G, Shearer WT. Immune function during space flight. Nutrition. 2002;18(10):899–903. doi:10.1016/s0899-9007(02)00903-6

Su M, Satola SW, Read TD. Genome-Based Prediction of Bacterial Antibiotic Resistance. J Clin Microbiol. 2019 Feb 27;57(3):e01405–18. doi: 10.1128/JCM.01405-18

Taylor PW. Impact of space flight on bacterial virulence and antibiotic susceptibility. Infect Drug Resist. 2015;8:249–262. Published 2015 Jul 30. doi:10.2147/IDR.S67275

Tixador R, Richoilley G, Gasset G, et al. Study of minimal inhibitory concentration of antibiotics on bacteria cultivated in vitro in space (Cytos 2 experiment). Aviat Space Environ Med. 1985;56(8):748–751

Urbaniak C, Sielaff AC, Frey KG, et al. Detection of antimicrobial resistance genes associated with the International Space Station environmental surfaces. Sci Rep. 2018;8(1):814. Published 2018 Jan 16. doi:10.1038/s41598-017-18506-4

Urbaniak C, Lorenzi H, Thissen J, et al. The influence of spaceflight on the astronaut salivary microbiome and the search for a microbiome biomarker for viral reactivation. Microbiome. 2020;8(1):56

Venkateswaran K, Vaishampayan P, Cisneros J, Pierson DL, Rogers SO, Perry J. International Space Station environmental microbiome - microbial inventories of ISS filter debris. Appl Microbiol Biotechnol. 2014;98(14):6453–6466. doi:10.1007/s00253-014-5650-6

Venkateswaran K, Singh NK, Checinska Sielaff A, et al. Non-Toxin-Producing Bacillus cereus Strains Belonging to the B. anthracis Clade Isolated from the International Space Station. mSystems. 2017;2(3):e00021–17. Published 2017 Jun 27. doi:10.1128/mSystems.00021-17

Voorhies AA, Mark Ott C, Mehta S, et al. Study of the impact of long-duration space missions at the International Space Station on the astronaut microbiome. Sci Rep. 2019;9(1):9911.

Wilson JW, Ott CM, Höner zu Bentrup K, et al. Space flight alters bacterial gene expression and virulence and reveals a role for global regulator Hfq. Proc Natl Acad Sci U S A. 2007;104(41):16299–16304. doi:10.1073/pnas.0707155104

World Health Organization. Global Action Plan on Antimicrobial Resistance (2015). Available online at: https://apps.who.int/iris/bitstream/handle/10665/193736/9789241509763_eng.pdf (accessed August 27, 2021).

Yoon S-H, Ha S-M, Lim J, Kwon S, Chun J. 2017. A large-scale evaluation of algorithms to calculate average nucleotide identity. Antonie Van Leeuwenhoek 110:1281–1286. https://doi.org/10.1007/s10482-017-0844-4.

Zankari E, Hasman H, Cosentino S et al. Identification of acquired antimicrobial resistance genes. J Antimicrob Chemother 2012; 67: 2640–4.

Zea L, Larsen M, Estante F, et al. Phenotypic Changes Exhibited by E. coli Cultured in Space. Front Microbiol. 2017;8:1598. Published 2017 Aug 28. doi:10.3389/fmicb.2017.01598

Zhang X et al. Fecal fusobacterium nucleatum for the diagnosis of colorectal tumor: a systematic review and meta-analysis. Cancer Med 8, 480–491 (2019)

