## Supplementary figures and images for "Machine learning algorithm to characterize antimicrobial resistance associated with the International Space Station surface microbiome"

### Supplementary Figure 1

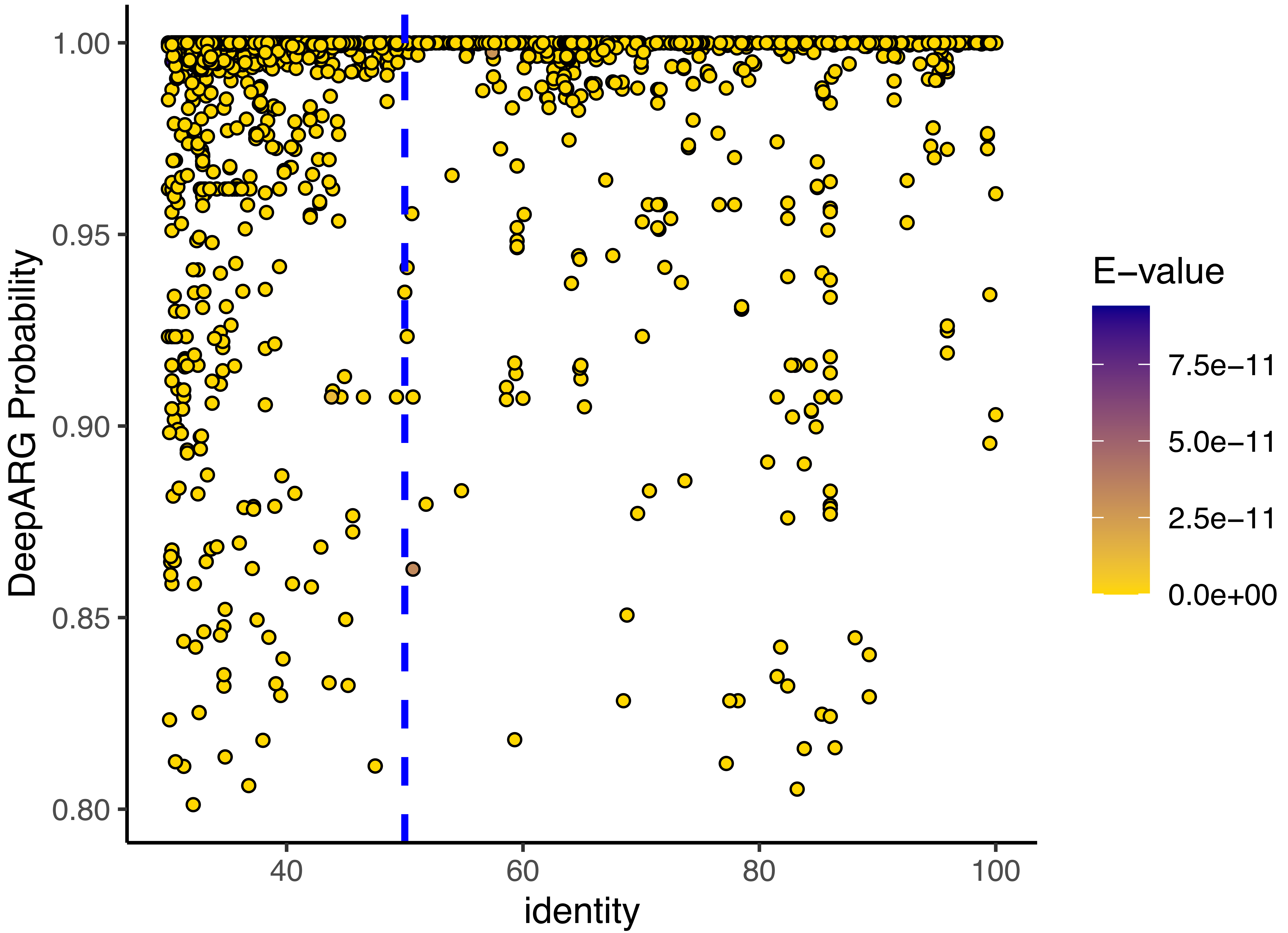
